# Androgen Receptor Signaling Regulates the Transcriptome of Prostate Cancer Cells by Modulating Global Alternative Splicing

**DOI:** 10.1101/828707

**Authors:** Kalpit Shah, Teresa Gagliano, Lisa Garland, Timothy O’Hanlon, Daria Bortolotti, Valentina Gentili, Roberta Rizzo, Georgios Giamas, Michael Dean

## Abstract

Androgen receptor (AR), is a transcription factor and a member of a hormone receptor superfamily. AR plays a vital role in the progression of prostate cancer and is a crucial target for therapeutic interventions. While the majority of advanced-stage prostate cancer patients will initially respond to the androgen-deprivation, the disease often progresses to castrate-resistant prostate cancer (CRPC). Interestingly, CRPC tumors continue to depend on hyperactive AR signaling and will respond to potent second-line anti-androgen therapies, including bicalutamide (CASODEX^®^) and enzalutamide (XTANDI^®^). However, the progression-free survival rate for the CRPC patients on anti-androgen therapies is only 8 to 19 months. Hence, there is a need to understand the mechanisms underlying CRPC progression and eventual treatment resistance. Here, we have leveraged next-generation sequencing and newly developed analytical methodologies to evaluate the role of AR-signaling in regulating the transcriptome of prostate cancer cells. The genomic and pharmacologic stimulation- and inhibition-of AR activity demonstrates that AR regulates alternative splicing within cancer-relevant genes. Furthermore, by integrating transcriptomic data from in vitro experiments and in prostate cancer patients, we found that a significant number of AR-regulated splicing events are associated with tumor progression. For example, we found evidence for an inadvertent AR-antagonist mediated switch in *IDH1* and *PL2G2A* isoform expression, which is associated with a decrease in overall survival of patients. Mechanistically, we discovered that the epithelial-specific splicing regulators (ESRP1 and ESRP2), flank many AR-regulated alternatively spliced exons. And, using 2D-invasion assays, we show that the inhibition of ESRPs can suppress AR-antagonist driven tumor invasion. In conclusion, until now, AR signaling has been primarily thought to modulate transcriptome of prostate epithelial cells by inducing or suppressing gene expression. Our work provides evidence for a new mechanism by which AR alters the transcriptome of prostate cancer cells by modulating alternative splicing. As such, our work has important implications for CRPC progression and development of resistance to treatment with bicalutamide and enzalutamide.

## Introduction

Androgen receptor (AR) is a member of the superfamily of hormonal nuclear receptors^1^. In the absence of its ligand, AR is secured in the cytoplasm by heat-shock proteins^2^. Once exposed to the male hormone androgen, AR, becomes activated, and translocates to the nucleus, where it binds to the androgen response elements (ARE) and initiate the transcriptional program^3-7^. Interestingly, activated AR molecules both enhance and suppress the expression of genes involved in prostate cancer progression^8-13^. This hormone-driven AR signaling is essential for development, differentiation, and normal functioning of the prostatic gland^14^. AR signaling, however, is hijacked in prostate tumors, driving disease progression. Therefore, the blockage of AR signaling through androgen deprivation continues to be the mainstay treatment of advanced-stage prostate cancer. While almost all patients with metastatic disease will initially respond to androgen ablation therapies, the majority of patients will progress to a castrate-resistant stage^15-18^.

Interestingly, studies employing xenograft prostate tumor models have shown that CRPC tumors that emerge after androgen-ablation therapy, continue to express AR and AR regulated genes^19^. Recent studies have argued that kinase-mediated hypersensitivity of AR^20-24^ and efficient uptake of androgens may play a critical role in fueling CRPC tumors^25^. Thus, the treatment option for patients with non-metastatic or metastatic CRPC typically includes high-affinity anti-androgens like bicalutamide (CASODEX^®^) and enzalutamide (XTANDI^®^)^26-28^. Although, in recent trials, enzalutamide has shown improved efficacy in comparison to bicalutamide the median time to PSA progression still suggests a limited benefit that lasts no more than 8 to 19 months^29^. In addition, in a few cases, an increase in metastasis of the disease was reported to be associated with the AR antagonist’s treatment regimen. Therefore, the search for the mechanism underlying CRPC, CRPC progression, and eventual treatment resistance would benefit patients who have exhausted all currently available treatment options. Towards this effort, the comprehensive understanding of AR functions continues to remain the center of focus.

The recent advent of high-throughput RNA sequencing and splicing microarrays has unveiled new layers of regulation of gene expression and highlighted the extreme complexity and versatility of the transcriptome. The majority of human genes encode multiple transcripts through the use of alternative promoters, alternative splicing (ASE), and alternative polyadenylation^30^. ASE is a mechanism that significantly expands the functional potential of the genome either by altering the usage of protein-coding transcripts, the ratio of coding to non-coding transcripts, or by allowing expression of isoforms with antagonistic functions from a single gene^31^. Multiple studies have found that ASE play a critical role in cancer^32^. A recent comprehensive analysis of alternative splicing across 32 cancer types from 8,705 patients revealed that tumors have up to 30% more alternative splicing events than normal tissues^33^. The steroid nuclear hormone receptors, including estrogen and progesterone receptors, are known to recruit regulators of ASE and modulate the transcriptome^34, 35^. However, whether modulation of AR-signaling may alter transcriptome of prostate cancer cells remains largely unexplored.

Herein, we hypothesized that modulation of AR-signaling either during prostate cancer progression or in response to treatment with AR antagonists might dysregulate the transcriptome of prostate cancer cells by modulating ASE. We employed a multitude of genomic approaches including Affymetrix splicing array, whole transcriptome RNA-seq analysis, and RT-PCR to show that AR-signaling regulates the transcriptome of prostate cancer cells by modulating ASE of a wide array of genes involved in regulating protein function. Furthermore, leveraging publicly available transcriptome data of primary-site samples from patients with prostate cancer at various stages of progression, we found a subset of AR-driven splicing events that are associated with progression of prostate cancer. Mechanistically, we found that Epithelial Splicing Regulator Proteins (ESRP1 & ESRP2) are the splicing factor through which AR may regulate splicing of pre-mRNA in prostate cancer cells. Interestingly, the inhibition of ESRPs suppressed AR-antagonist mediated increase in the invasion of prostate cancer cells. Collectively, we provide the evidence for a novel and critical mechanism of prostate cancer progression that is regulated by AR and that the treatment with AR-antagonist may inadvertently promote invasion by dysregulating splicing of critical genes.

### Pharmacological Manipulation of Androgen Receptor Signaling Induces Alternative Splicing in Prostate Cancer Cells

To study the effect of pharmacological manipulation of AR signaling in prostate cancer cells, we performed expression profiling of LNCaP cells that were cultured for 72 hours in charcoal-stripped fetal bovine serum and stimulated with 10nM AR agonist, dihydrotestosterone (DHT), or 10uM of the AR antagonist casodex for 24 hours. The array consisted of > 6 million probes, of which 70% covered exons for coding transcripts while the remaining 30% covered exon-exon splice junctions and non-coding transcripts; hence, allowing us to monitor transcriptional changes at the level of the exon. Besides, the well-characterized gene expression changes (Supplementary Table 1), we found that pharmacological manipulation of AR signaling induced global changes in ASE, which was evident by differential expression of exon-exon splice junction probes in comparison to constitutive exons (Supplementary Table 2). Figure 1A shows the expression of top-50 differentially expressed probes spanning exon-exon junction of a gene across different conditions. We next sought to characterize the potential ASE events using the transcription analysis console, which integrates the evidences from array probes spanning exon-exon splice junction and constitutive exons to classify the events as either cassette exon (CE), alternative 3 prime start site (A3SS), alternative 5 prime start site (A5SS), intron retention (IR), complex event, alternative last exon (ALE), mutually exclusive exon (MEE), or alternative first exon (AFE) (Figure 1B). Because changes in gene expression may confound ASE call, we filtered out any splicing events that occurred within gene that were differentially expressed in treatment group in comparison to control. In summary, treatment of LNCaP cells with DHT or casodex resulted in 671 and 2127 significant ASE events in comparison to DMSO treatment respectively. Furthermore, in comparison to stimulation, inhibition of AR in LNCaP cells led to greater than 2827 ASE events. We found that treatment of LNCaP cells with antagonist or agonist did not drastically alter the distribution of CE (63% vs 84%), A3SS (12% vs 6%), A5SS (15.0% vs 8.0%), ALE (0.2% vs 0%), MEE (0% vs 0.1%), and AFE (0.1% vs 0%). However, treatment with casodex did show a ten-fold increase (11.0% vs 1.0%) in the percentage of IR events in comparison to agonist treated LNCaP cells.

**Figure 1:**
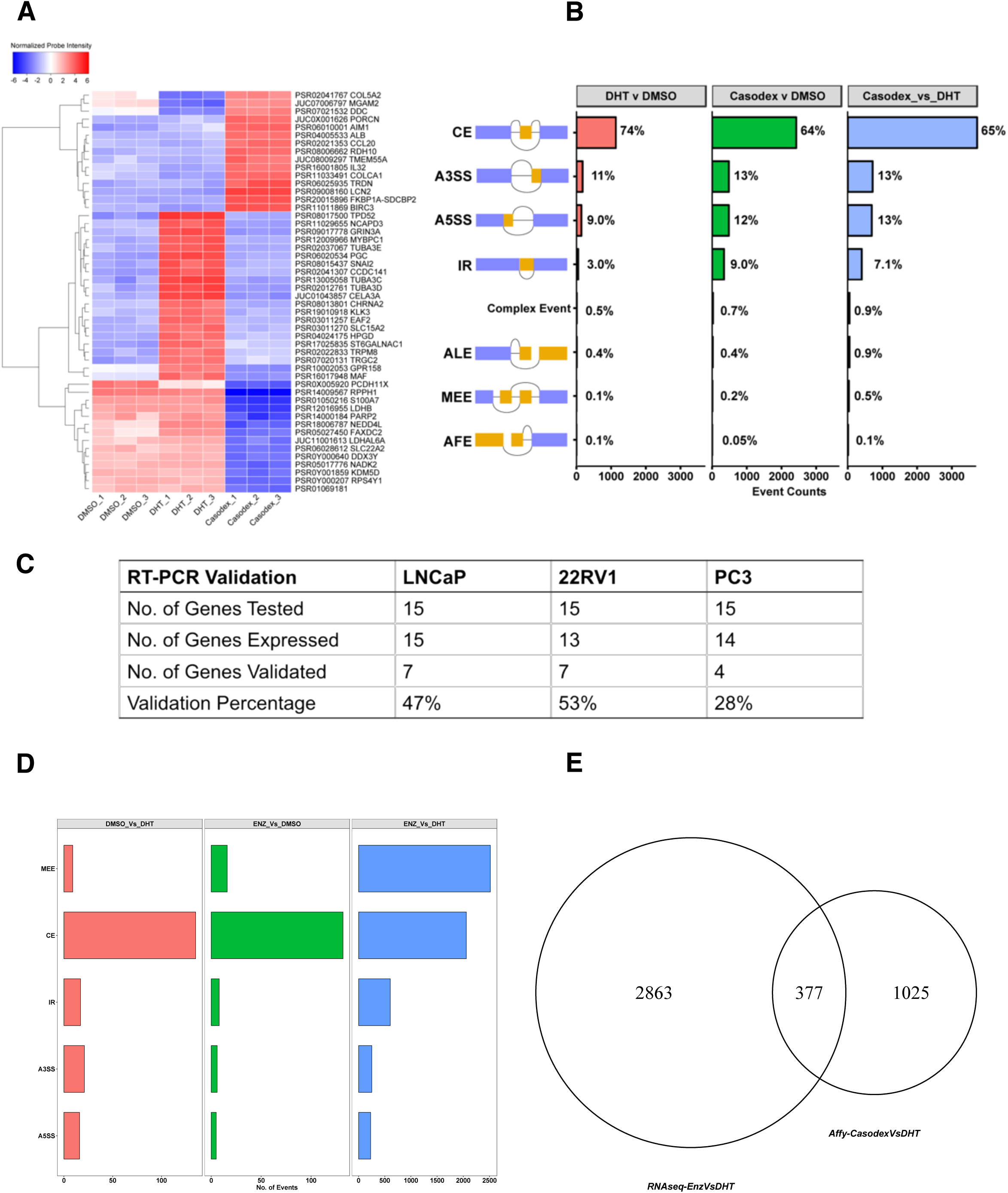
Pharmacological Inhibition of Androgen Receptor Signaling Induces Alternative Splicing in Prostate Cancer Cells. (A) Heatmap showing the normalized probe intensity of top-50 differentially expressed probes spanning exon-exon junction of a gene across different conditions including LNCaP cells cultured for three days in CSFBS and treated with either vehicle (DMSO), 10nM DHT, or 10μM casodex. (B) Bar plot detailing percentage of alternative splicing events including CE, A3SS, A5SS, IR, ALE, MEE, AFE, and complex events in LNCaP cells across three comparisons including 10nMDHT vs DMSO, 10μM casodex vs DMSO, and 10μM casodex vs 10nMDHT. (C) We leveraged RT-PCR and validated 15 splicing events, which were rationally selected from the affy transcriptomic analysis in three prostate cancer cell line models. Briefly LNCaP, 22RV1, and PC3 cells were cultured in charcoal stripped fetal bovine serum for 3 days and treated with either DMSO, 10nM DHT, or 10μM casodex. The table details no. of genes tested, no. of genes expressed in each cell line, and no. of genes that were validated using RT-PCR assay. (D) Bar plot detailing percentage of alternative splicing events including CE, A3SS, A5SS, MEE and IR in LNCaP cells across three comparisons including DMSO vs DHT, enzalutamide vs DMSO, and enzalutamide vs DHT. (E) Venn diagram comparing genes identified to undergo ASE in LNCaP cells treated with either casodex or enzalutamide.

We next sought to validate the ASE events predicted by the splicing array using RT-PCR assay. The CE and IR are some of the most common splicing events contributing towards transcriptional heterogeneity in tumor cells and are also well characterized^33, 36-38^. Therefore, we decided to validate a total of 15 of these events in three separate cell lines including LNCaP, 22RV1 (castrate-resistant prostate adenocarcinoma cells), and PC3 (bone metastatic prostate cancer cells). We performed three independent experiment each with three technical replicates. The ASE events for validation were picked based on the following three criteria 1) Events with FDR cut off of 0.05 and SI of >= |2|; 2) Events with evidence from not only the probes mapping to the alternatively spliced region but also those that mapped to junction surrounding it, and 3) Genes with a known biological role in cancer. We used two primer pairs, one for monitoring the expression of constitutive exons within all the isoforms of a gene and another for measuring changes in the alternatively spliced region. The primers specific to the constitutive exons revealed that LNCaP cells express 15 genes, 22RV1 express 13, and PC3 cells express 14 of 15 genes tested (Supplementary Table 3). The RT-PCR assay validated 47%, 53%, and 28% of HTA-2.0 predicted splicing events in LNCaP, 22RV1, and PC3 cells respectively (Figure 1C and Supplementary Figure 1–3, Supplementary Table 4).

We next leveraged publicly available and in-house generated whole transcriptomic data to investigate whether treatment with enzalutamide, a more potent AR antagonist than casodex would also induce ASE in LNCaP cells. Briefly, we performed strand-specific 150-bp paired-end RNA-seq for LNCaP cells treated with vehicle or 10nM DHT. Additionally, we downloaded a dataset for enzalutamide or vehicle treated LNCaP cells from GSE110903. Altogether, we compiled data consisting of two biological replicates per sample with 45–80 million mapped reads per replicate. We used the rMATS computational pipeline with default settings to identify the splicing changes. The analysis revealed that DHT and enzalutamide treatment induced 198 and 167 significant ASE events respectively at a stringent filter of FDR <= 0.05 and delta PSI of >= 10%. The largest difference in ASE (∼5663) was observed when we compared the transcriptome of LNCaP treated with enzalutamide with that of DHT (Figure 1D). Furthermore, rMATS classification of enzalutamide or DHT induced ASE revealed that the majority of splicing events were either MEE or CE. The heatmap showing the top 100 significantly different PSI for MEE and CE across LNCaP cells treated with enzalutamide or DHT is shown in Supplementary Figure 4A. Moreover, splicing analysis using whole transcriptome and splicing array data are reported to produce discordant results and consistent with that observation we too found a very small overlap (377/4265) between genes identified to undergo ASE upon treatment with casodex or enzalutamide (Figure 1E). Overall, evidence from analysis of splice array, RT-PCR and whole transcriptome data supports our novel observation that modulation of AR signaling alters transcriptome of prostate cancer cells by regulating ASE.

### Functional Analysis of Genes Regulated at the Level of Alternative Splicing and Transcription in Prostate Cancer Cells

In order to study the significance of ASE in prostate cancer cells that are driven by pharmacological manipulation of AR signaling, we first queried whether treatment with enzalutamide or casodex altered splicing of prostate cancer relevant genes. We curated a list of 100 genes (Supplementary Table 5) that are associated with prostate cancer progression and development. Our analysis showed that casodex induced ASE in 49 prostate cancer relevant genes including *KLK3, AKT2, EGFR, PIK3R, KRAS, VDR*, and *MTOR*. Similarly, enzalutamide induced ASE in 19 prostate cancer relevant genes. The heatmap comparing PSI of the prostate cancer relevant genes across samples is shown in Figure 2A. One of the RT-PCR validated ASE prostate cancer genes is *IDH1*, a key metabolic gene regulating TET2 mediated epigenetic re-programing in prostate tumor cells^39, 40^. We found that casodex treatment of all three prostate cancer cell lines resulted in a switch from ENST00000345146, a dominant transcript of IDH1 to ENST00000415913 with an alternate 5’UTR (Supplementary Figure 1). The translational relevance of this functional switch was accentuated by our expression and survival analyses, which revealed that the expression of the primary isoform (ENST00000345146) is significantly higher (p = 1.84e-43) in the TCGA-Prostate Adenocarcinoma tissue (N = 495) in comparison to the GTEX-normal prostate tissue (N = 100) and is also associated with decreased overall patient survival (Figure 2B).

**Figure 2:**
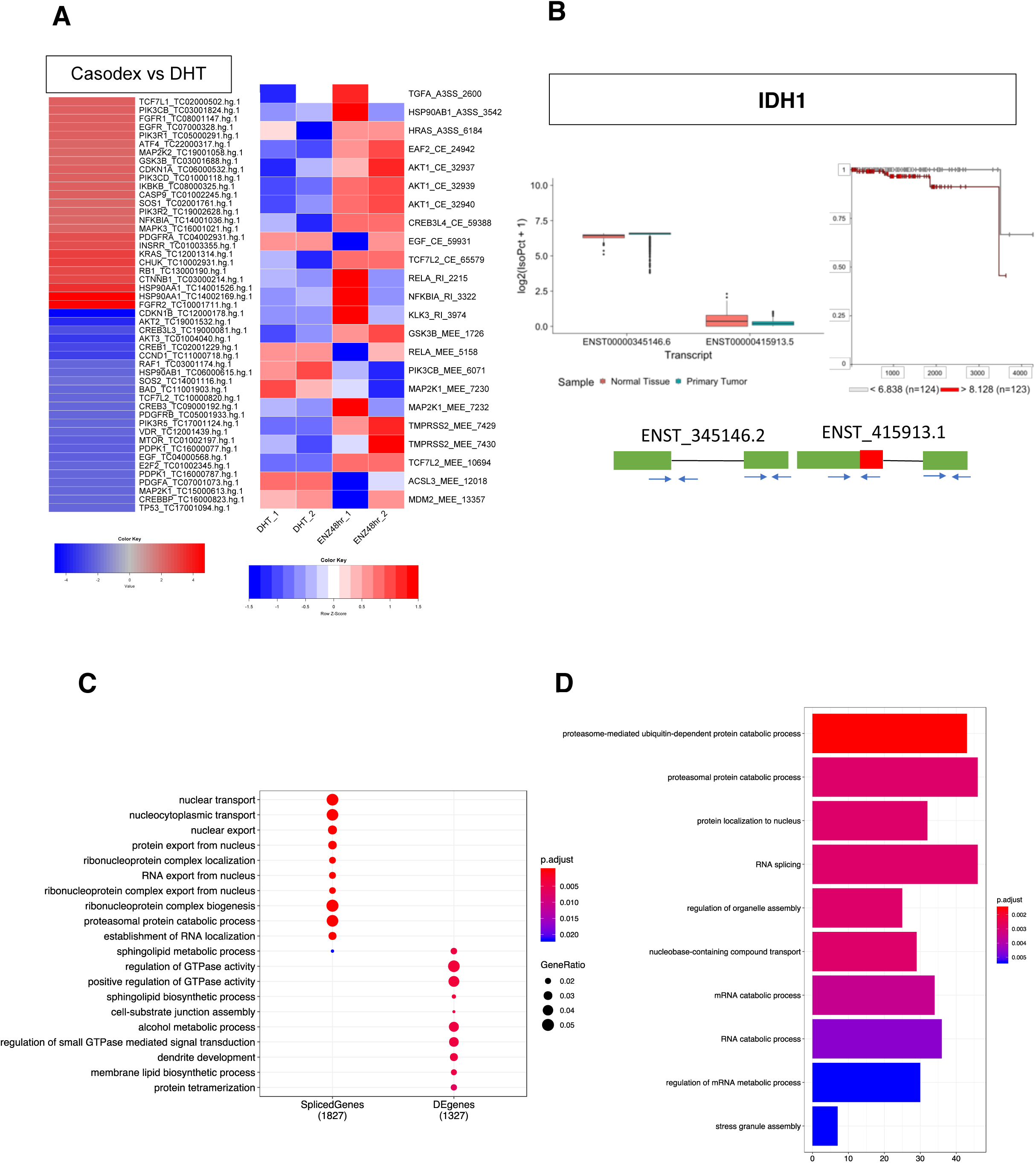
Functional Analysis of Genes Regulated at the Level of Alternative Splicing and Transcription in Prostate Cancer Cells. (A) Heatmap showing the differential percent splice index (PSI) of prostate cancer relevant genes between LNCaP cells cultured with 10μM Casodex and 10nM DHT. The adjacent heatmap shows the PSI across different conditions including LNCaP cells treated with DHT or enzalutamide. (B) Functional validation for the Casodex mediated switch in the IDH1 isoform expression. The box plot comparing the expression of the ENST00000345146 or ENST00000415913 between TCGA-prostate adenocarcinoma tissue and GTEX-normal prostate tissue. The kaplan-meier plot displays the association between expression of ENST00000345146 and survival for patients with prostate cancer. (C) A dot-plot comparing GO pathways enriched in differentially expressed or alternatively spliced genes modulated by casodex and DHT. (D) A bar plot showing GO pathways enriched in genes modulated by enzalutamide in comparison to DHT.

We further investigated whether genes modulated by casodex and DHT at the level of transcription and ASE have different physiological roles. We derived biological roles for this exclusive set of AR-axis modulated genes by using gene ontology (GO) overrepresentation analysis. We used all the genes that have an annotation as reference in our analysis as recommended by clusterProfiler. The alternatively spliced genes were enriched in pathways related to modulation of gene expression including nucleic acid, protein transport, mRNA splicing, and proteosomal degradation whereas the differentially expressed genes were enriched in pathway related to EMT including those involved in GTPase activity, cell-cell interaction/junctions, and cytoskeleton organization (Figure 2C). We next leveraged RNAseq data from enzalutamide and DHT treated cells to investigate the potential mechanisms of action through which ASE may alter function of a gene. Briefly, we mapped the region undergoing ASE to protein domains of the gene using Maser R package. We found that the majority of splicing occurred in the characterized functional domains including the transmembrane domain, coiled region, topo domain, metal binding, zinc finger binding, and activation site for protein (Supplementary Figure 4B). Furthermore, similar to casodex, enzalutamide treatment of LNCaP cells also dysregulated splicing of genes enriched in pathways involved in the modulation of gene expression including splicing, transport of nucleic acid, proteasomal degradation, and protein localization (Figure 2D). Collectively, our results strongly suggest that the changes in androgen-driven ASE are biologically meaningful and distinct from the functional impact of androgen-driven gene expression changes.

### Direct Genomic Inhibition of Androgen Receptor in Prostate Cancer Cells Induces Alternative Splicing

Our data provides evidence for the AR agonist and antagonist mediated regulation of ASE of pre-mRNA. However, it is possible that the observed ASE is a non-specific effect from the pharmacological treatment of cells and not a direct effect by modulation of AR. To test this possibility, we modulated expression of AR in 22RV1 cells using siRNA and used RT-PCR assays to study altered splicing of genes including *AAK1, SYNE4, and MAN1A1* which were predicted to undergo ASE in response to casodex treatment. We found a robust five-fold decrease in the expression of AR in 22RV1 cells transfected with siRNA in comparison to a scrambled siRNA (Figure 3A). Supporting our hypothesis, we found that the five-fold inhibition of AR altered splicing of *AAK1* (SI: −2.52), *SYNE4* (SI: − 1.36), and *MAN1A1* (SI: −3.22) in the same direction as that of casodex (Figure 3B, Supplementary Table 4). This data supports our hypothesis that ASE events induced by agonists or antagonists of AR are driven in part by direct modulation of AR.

**Figure 3:**
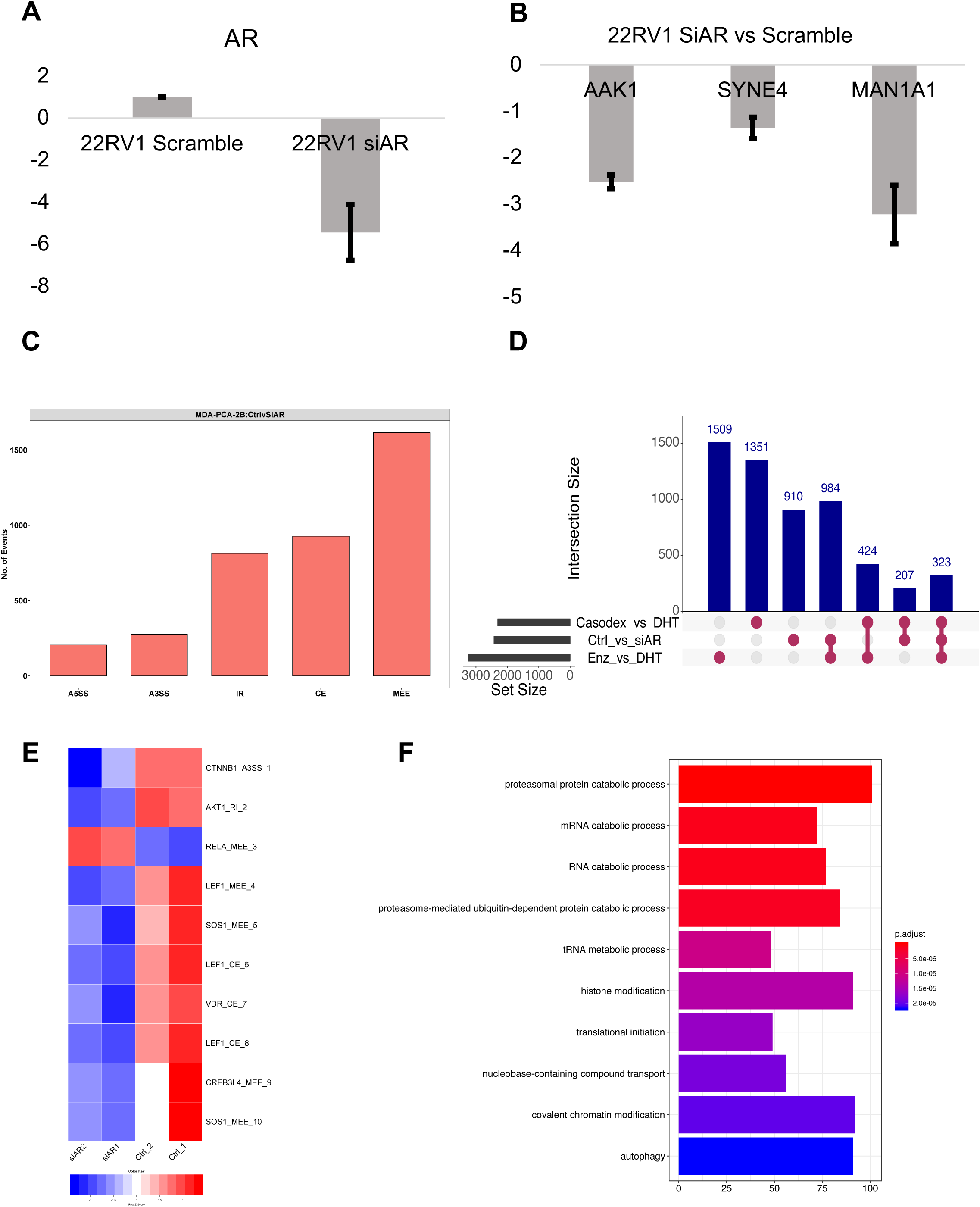
Direct Genomic Inhibition of Androgen Receptor in Prostate Cancer Cells Induces Alternative Splicing. (A) Bar-graph comparing expression of AR in 22RV1 treated with scramble siRNA or siRNA against AR. (B) Bar-graph showing expression of AAK1, SYNE4, and MAN1A1 in LNCaP cells treated with siRNA against AR in comparison to scramble siRNA. (C) Bar-graph showing total number of rMATS predicted splicing events in MDA-PCa-2b cells treated with siRNA against AR or scrambled control. (D) Up-set plot comparing the genes predicted to undergo alternative splicing by rMATS or HTA2.0 analysis in prostate cancer cells treated with casodex or enzalutamide in comparison to DHT or siRNA against AR in comparison to scrambled siRNA. (E) The heatmap comparing PSI for prostate cancer genes across MDA-PCa-2b cells treated with scrambled or siRNA against AR. (F) Bar-graph revealing the GO pathway enriched for genes under undergoing ASE modulated by genomic inhibition of AR in MDA-PCa-2b.

We next leveraged publicly available (GSE110903) RNAseq data to further study the effects of genomic inhibition of AR on ASE in MDA-PCa-2b cells, a model for advanced prostate cancer bone metastasis cells that express PSA, AR and are androgen sensitive. The rMATS analysis revealed that siRNA knockdown of AR in MDA-PCa-2b cells induced 3841 ASE events after a stringent filtration of FDR <= 0.05 and delta PSI of >= 10%. Also, similar to our observations with enzalutamide treated LNCaP cells, we found that the MEE formed the largest fraction of ASE events induced by the siRNA knockdown in MDA-PCa-2b and were followed by CE and IR events (Figure 3C). In addition, we used the upset plot for identifying the overlap between genes that are regulated at the level of splicing by the pharmacological inhibitors of AR in LNCaP cells or genomic inhibition of AR in MDA-PCa-2b. We found an overlap of 984 genes between enzalutamide treated LNCaP and siRNA treated MDA-PCa-2b cells, 424 genes between casodex and enzalutamide treated LNCaP cells, and 207 genes between casodex treated LNCaP and siRNA treated MDA-PCa-2b cells. Interestingly, 323 genes were common between all three treatment groups and were enriched for pathways involved in the regulation of translation (Figure 3D, Supplementary Figure 4C). The plot also revealed a non-overlapping set of genes in all treatment groups, possibly indicating a combination of differences in prostate cell line models and assays used for measuring ASE. The ASE genes also included known prostate cancer relevant genes including *CTNNB1, AKT1, LEF1*, and *VDR*. The heatmap comparing PSI for prostate cancer genes across MDA-PCa-2b cells treated with scrambled or siRNA against AR (Figure 3E). Similar to enzalutamide treated LNCaP cells, the GO pathway analyses revealed that genes undergoing ASE modulated by genomic inhibition of AR are enriched for pathways including mRNA splicing, translation initiation, chromatin remodeling, epigenetic regulation, and proteasomal degradation (Figure 3F). In addition, the functional mapping of alternatively spliced exons revealed that genomic inhibition of AR may dysregulate function of the alternatively spliced genes by modifying functional domains (Supplementary Figure 4D). Overall, we provide two lines of evidences supporting our hypothesis that manipulation of AR-axis either by pharmacological inhibitors of AR or by siRNA alters the splicing of pre-mRNA, modifies the functional domain of gene and consequentially dysregulates its function.

### Androgen Receptor-Axis Changes Splicing of pre-mRNA by Modulating Expression of ESRP1/ESRP2, the Master Regulator of Alternative Splicing

RNA binding proteins (RBPs) are key proteins that bind to mRNA or non-coding RNAs and play a wide variety of roles in post-transcriptional processing including regulation of ASE. Hence, we hypothesized that RBPs which are differentially expressed in response to modulation of AR-axis might provide mechanistic insight into the observed regulation of ASE. To test our hypothesis, we curated a list of 112 RBPs from the published literature^41^ with a known role in splicing regulation and investigated if their expression was regulated by pharmacological manipulation of AR. Interestingly, we found that out of 112 RBPs only Epithelial Splicing Regulator Proteins (ESRP1/2) were dysregulated by casodex and DHT (Figure 4A, & Supplementary Table 1). Furthermore, numerous studies have also reported that splicing is enhanced when ESRP1/2 binds to pre-mRNA downstream of the exon, while splicing is enhanced when ESRP1/2 binds upstream of or within the exon. Therefore, to gain additional support for the interplay between ESRP1/2 and AR, we performed spatial analysis of ESRP1/2 binding sites around the exons that are alternatively spliced in response to pharmacological inhibition of AR in the LNCaP cell line. We performed a spatial analysis for all the significant cassette exon events in LNCaP cells treated with enzalutamide in comparison to DHT. Figure 4B shows that there is enrichment of ESRP2 binding sites in the intronic region 0 to 125nt downstream of AR-upregulated exons and underrepresented in the same region downstream of AR-downregulated exons. In contrast, we found that there is underrepresentation of ESRP1 binding sites in the intronic region 0 to 125nt downstream of AR-upregulated exon and enrichment in the same region downstream of AR-downregulated exons. Furthermore, ESRP2 binding motifs were enriched upstream of AR-downregulated exons between −50 and −150 and underrepresented in the same region of AR-upregulated exons. In contrast, ESRP1 binding motif were enriched upstream of the AR-upregulated exons between 0 and −100 and underepresented in the same region of AR-downregulated exons. In addition, while ESRP2 motifs were under-represented in the region within the silenced and upregulated exons, ESRP1 motif were enriched in the upregulated exon and under-represented in downregulated exons. This suggest that pharmacological manipulation of AR may alter splicing by regulating expression and binding of ESRP1/2 around spliced exons.

**Figure 4:**
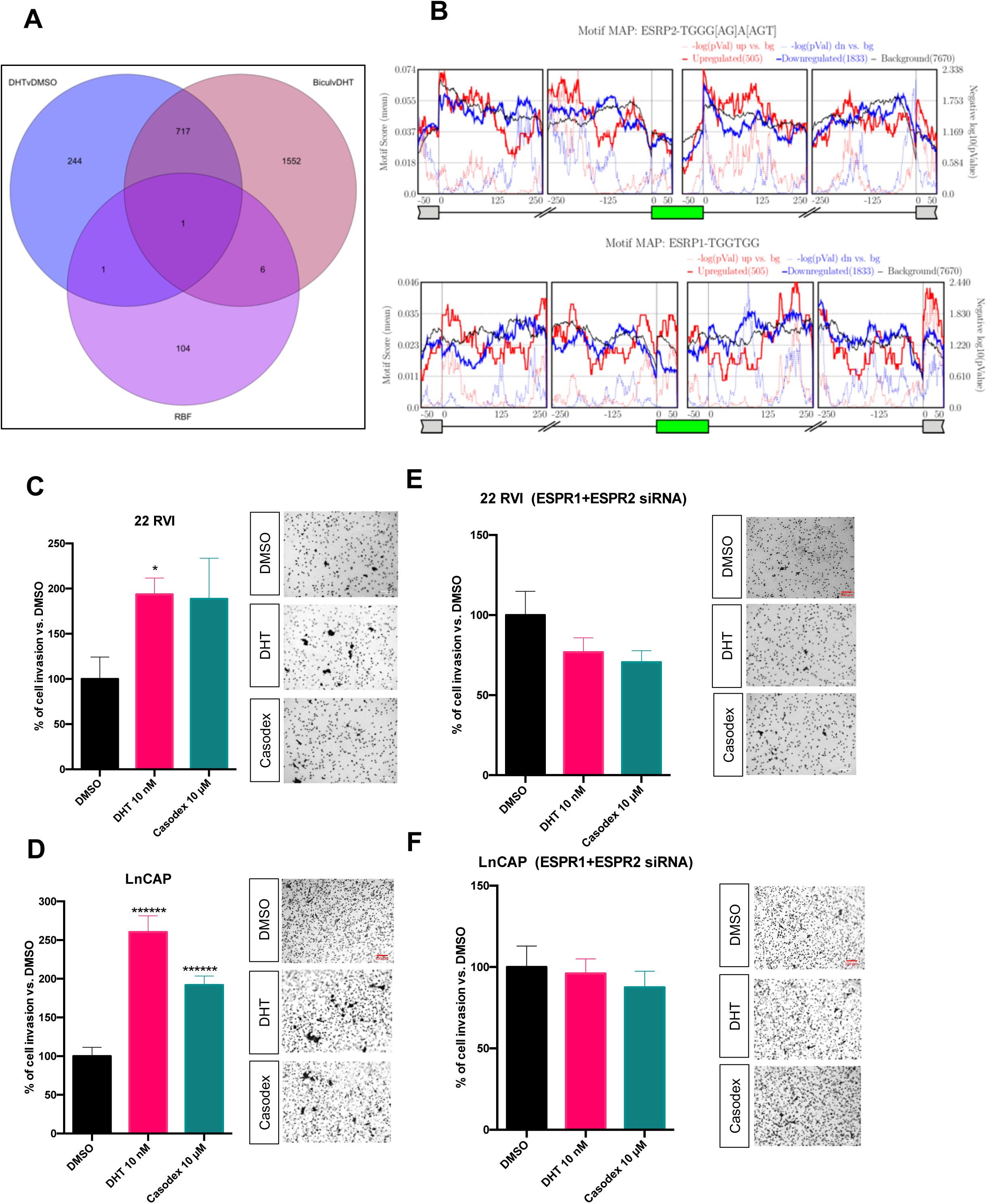
Androgen Receptor-Axis Changes Splicing of pre-mRNA by Modulating Expression of ESRP1/ESRP2, the Master Regulator of Alternative Splicing. (A) Venn-diagram showing an overlap between a curated list of 112 RBPs with known role in regulating alternative splicing, genes that were differentially expressed in LNCaP cells treated with DHT or Casodex in comparison to DMSO. (B) Maps for ESRP1 and ESRP2 binding motifs showing enrichment upstream and downstream from exons upregulated or downregulated in LNCaP cells treated with enzalutamide in comparison to DHT. 2D invasion assay: 22 RVI (C) and LnCAP (D) cells were seeded on the Matrigel-coated upper chamber of the transwell insert and treated with either vehicle (DMSO) or with 10 nM DHT or 10 μM Casodex in serum free media, while 10% FBS medium was added to the lower chamber used as chemoattractant. After 24h migrated cells were fixed and stained with crystal violet and counted using an inverted microscope. One-way ANOVA test was performed using Prism 8 software. Data are expressed as mean ± SEM; * P < 0.05, ****** P < 0.00001. Experiment was performed 3 time, with 3 replicates for each experiment. 22 RVI (E) and LnCAP (F) cells were transfected with indicated siRNA cell, seeded on the Matrigel-coated upper chamber of the transwell insert and treated with either vehicle (DMSO) or with 10 nM DHT or 10 μM Casodex in serum free media, while 10% FBS medium was added to the lower chamber used as chemoattractant. After 24h migrated cells were fixed and stained with crystal violet and counted using an inverted microscope. One-way ANOVA test was performed using Prism 8 software. Data are expressed as mean ± SEM; no significand was detected. Experiment was performed 3 time, with 3 replicates for each experiment.

Furthermore, ESRP1/2 are involved in EMT and invasion of cancer cells^42, 43^. However, the role of ESRP1/2 in prostate cancer has not been established. In order to study the significance of the AR-ESRP axis in prostate cancer, we employed an in vitro invasion assay. Consistent with published *in vitro* and *in vivo* report^44^, we found that treatment with DHT and casodex increases the invasion rate of LNCAP and 22RV1 (Figure 4C & D). Interestingly, when ESRP1/2 is silenced in prostate cancer cells (Figure 4E & F), the DHT or casodex mediated increase in the invasion rate is completely abolished. Furthermore, the genomic inhibition of ESRP1/2 in prostate cancer cells did not affect the expression of E-cadherin and vimentin, key genes involved in EMT (Supplementary Figure 6).

### Modulation of AR-axis regulates splicing of pre-mRNA that are associated with progression of prostate cancer disease

Our analysis in prostate cancer cell lines shows that modulation of AR signaling dysregulates splicing of functionally relevant genes. Since dysregulation of AR signaling is the hallmark of prostate cancer progression, we hypothesized that ASE of functionally relevant genes would be associated with the progression of disease. To test our hypothesis, we employed rMATS splicing analysis using GSE80609^45^. This dataset consisted of 8 benign prostate hyperplasia (BPH), 16 localized prostate cancer (L.PC), nine advanced prostate cancer (advPC), 12 castrate-resistant prostate cancer (CRPC), and four pairs of advPC and CRPC samples from the same patient. Since the rate for RT-PCR validation for bioinformatically predicted spliced events is low, we filtered our results with stringent cut-off of at least 10% difference in PSI and minimum FDR value of at least 0.05. In concordance with Kang et al. gene-centric study, our splice analysis also found the greatest difference between the transcriptome of BPH and L.PC and lowest between that of advPC and CRPC (Figure 5A). In particular, we found that 53 A3SS, 46 A5SS, 574 CE, 191 IR, and 277 MEE differentiated BPH from L.PC (A); 25 A3SS, 13 A5SS, 155 CE, 81 IR, and 128 MEE events differentiated L.PC from adv.PC (B); 8 A3SS, 10 A5SS, 73 CE, 20 IR, and 14 MEE differentiated adv.PC from CRPC (C); and 10 A3SS, 20 A5SS, 72 CE, 37 IR, and 2 MEE differentiated paired samples of advPC and CRPC (D) (Figure 5A).

**Figure 5:**
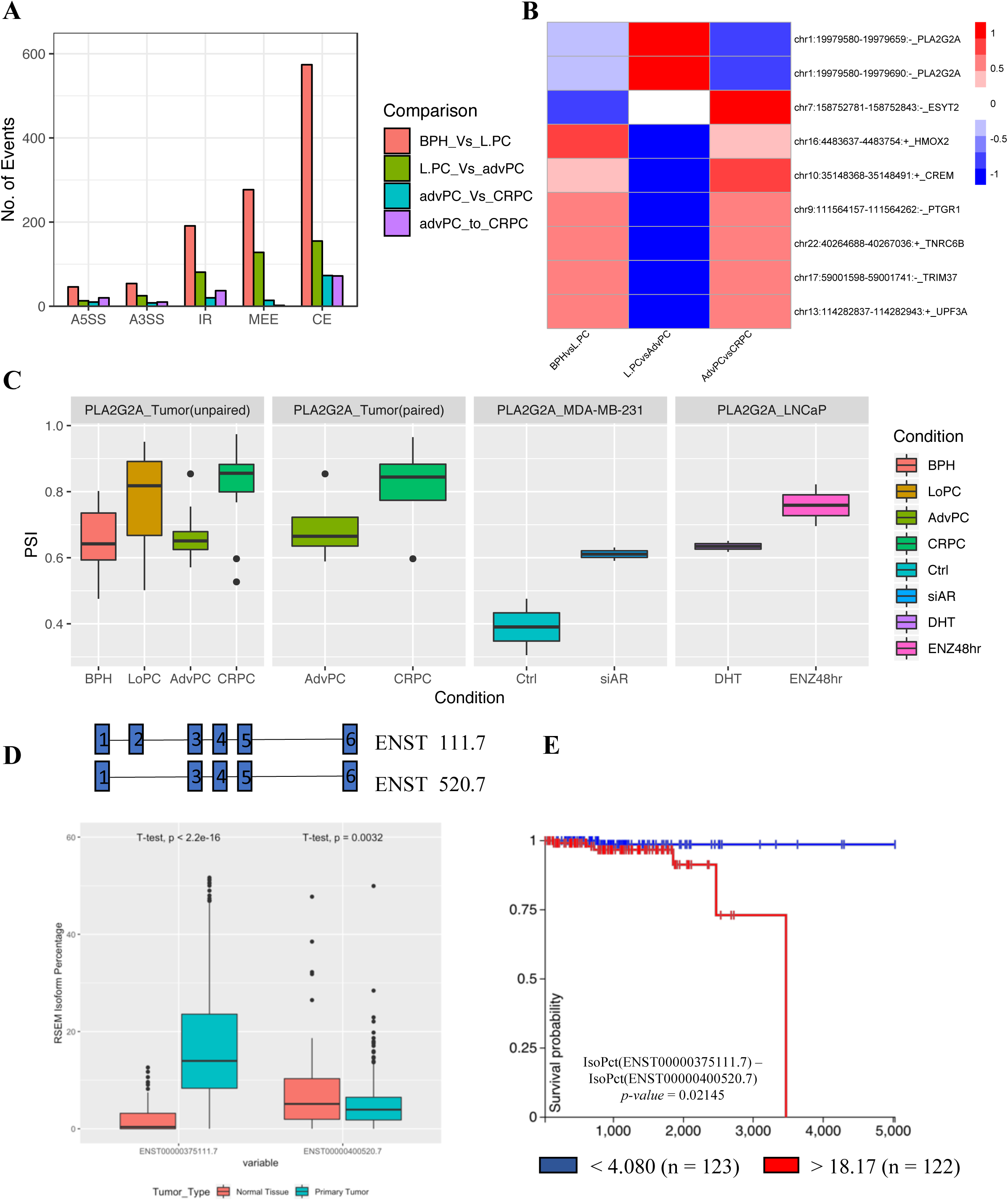
Modulation of AR-axis regulates splicing of pre-mRNA that are associated with progression of prostate cancer disease. (A) Bar-graph comparing significant number of splicing events across patients at different stages of prostate cancer including benign prostate hyperplasia (BPH), localized prostate cancer (L.PC), advanced prostate cancer (advPC), castrate-resistant prostate cancer (CRPC), and four pairs of advPC and CRPC samples from the same patient. (B) Heatmap comparing the PSI for the cassette or mutually exclusive exons across patients at different stages of prostate cancer. (C) Box plot displaying the changes in PSI of the exon-2 of PLA2G2A gene in patients at different stages of prostate cancer or in prostate cancer cells treated with DMSO, DHT, Enzalutamide, scrambled siRNA, or siRNA against AR. (D) The box plot comparing the expression of the ENST00000375111.7 or ENST00000400520.7 between TCGA-prostate adenocarcinoma tissue and GTEX-normal prostate tissue. (E) The kaplan-meier plot displays the association between overall survival for patients with prostate cancer and difference in expression of percentage isoform of ENST00000375111.7 than ENST00000400520.7.

The earlier study identified dysregulation of AR expression as the only shared event across all stages of prostate cancer^45^. To the contrary, we found a total of 9 splice events to be associated with all stages of prostate cancer (A⋂B⋂C⋂D) (Supplementary Figure 5A–C). These events included CE and MEE in the pre-mRNA of cancer-relevant genes, including *TRIM37, PTGR1, CREM, HMOX2, UPF3A, TNRC6B, PLA2G2A, and ESYT2*. The direction of PSI for these events varied during disease progression, suggesting a differential role for these genes in disease (Figure 5B). Supplementary Figure 5D-K shows the transcript plots displaying the usage of exon for these nine genes between AdvPC and CRPC. The transcriptome of advPC and CRPC is reported to be highly similar and previous study had identified only 15 genes that differentiated their transcriptome. Hence, we further investigated whether ASE events may further distinguish advPC from CRPC. We compared the list of splicing events in paired and unpaired samples of advPC and CRPC (C⋂D) and found 13 unique splicing events that differentiated the transcriptome of advPC and CRPC (Supplementary Figure 5A–C). These splicing events occurred in the pre-mRNA of *PTGR1, FRG1HP, RP11, CA5BP1, CREM, TNRC6B, BCS1L, FASTKD1, ESYT2, CLN3, PLA2G2A, MYL6, FBXL12, ZNF202, UBAP2, and MIR940* (Supplementary Figure 5L). The rMATS analysis of the prostate cancer dataset identified that ASE is associated with tumorigenesis, advanced progression, and CRPC development. Furthermore, in addition to differentially expressed genes, our study identifies splicing events that can further differentiate the transcriptome of advPC and CRPC.

Our analysis shows that pharmacological or genomic inhibition of AR alters splicing of several functionally relevant genes. Therefore, it is possible that a subset of these alternatively spliced genes is associated with progression of prostate cancer. To test this possibility, we compared the list of splicing events common across all stages of prostate cancer (A⋂B⋂C⋂D) and list of splicing events induced by either pharmacological and genomic inhibition of AR in LNCaP and MDA-PCa-2b cells (Supplementary Figure 5A– C). We found only one significant event that alters the inclusion of exon-2 (Chr1:19979580-19979659) in the pre-mRNA of *PLA2G2A*, a secreted phospholipase. PLA2G2A is significantly downregulated in patients with metastatic prostate cancer in comparison to the primary tumor. Moreover, the decrease in expression of *PLA2G2A* is implicated in promoting invasion and metastasis^46^. Interestingly, exon-2 of *PLA2G2A* contains a repressor region, and its inclusion is associated with a decrease in the expression of the gene^47, 48^. In support of the known role of *PLA2G2A*, we found an increase in the inclusion of exon-2 in advPC in comparison to CRPC (paired and unpaired samples), BPH in comparison to L.PC (unpaired samples) and in response to the pharmacological or genomic inhibition of AR signaling in LNCaP and MDA-PCa-2b cell lines respectively (Figure 5C).

To study the functional significance of the exon-2 splicing, we performed expression and survival analysis with the TCGA–PRAD and GTEX datasets. In particular, we found that percentage of the ENST00000375111.7 isoform, that contains exon 2 was significantly higher in patients with prostate cancer (n = 496, TCGA-PRAD) in comparison to healthy prostate tissue (n = 100, GTEX-prostate). In contrast, the percentage of the ENST00000400520.7 isoform, that does not contain exon 2 was significantly lower in patients with prostate cancer in comparison to healthy prostate (Figure 5D). In addition, the survival analysis for primary prostate cancer patients in TCGA dataset revealed that patients with primary tumors with a higher difference between percentage isoform of ENST00000375111.7 than ENST00000400520.7 (> 18.17; n = 122) had significantly (p = 0.021) shorter overall survival (OS) compared to those difference of < 4.08 (n = 123) (Figure 5E). In conclusion, we show that differential usage of exon or introns is associated with progression of prostate cancer. In addition, inhibition of AR may inadvertently switch the delicate balance between different splice isoforms in genes critical for disease progression.

## DISCUSSION

Prostate cancer is driven by dysregulation of transcriptional networks regulated by molecules, including AR, FOXA1, and ERG. The overwhelming evidence in the field points to the fact that of all the transcription factors, AR not only plays a critical role in initiation but also, the development and progression of prostate cancer. The role of AR in prostate cancer is signified by the fact that the current mainstay therapies for prostate cancer involve modulating AR-transcriptional activity by chemical castration. Castration therapy works well during the initial stages of tumor development; however, the disease often relapses as CRPC. Because CRPC tumors continue to depend on AR-transcriptional activity, these patients will respond to second-line anti-androgen therapies, including bicalutamide and enzalutamide. However, CRPC patients on anti-androgen therapies will only experience 8 to 19 months of progression-free survival^29^. Hence, it is critical to understand how AR regulates gene transcription in healthy and disease state.

Alternative splicing (ASE) is known to play a significant role in maintaining cellular physiology, and the identification of transcriptional splicing patterns may have the potential for early diagnosis, prognosis, and identification of therapeutic targets in tumor biology^49^. The nuclear hormone receptors, including estrogen and progesterone receptors, are known to modulate ASE. However, the role of AR in modulating ASE remains unexplored. Therefore, in this study, we focused on the role of AR, AR-signaling, and its therapeutic inhibitors in regulating ASE in prostate cancer and its implication on tumor physiology and progression. Earlier work used a targeted approach with minigenes to discover the role of nuclear receptors in modulating splicing. We leveraged HTA2.0, a newer-generation of microarray that interrogates junctions between exons in the transcriptome as well as the exon themselves and RNA-Seq analysis to test whether modulating AR-signaling would alter ASE prostate cancer cells. This approach led to an unexpected finding that treatment with DHT or Casodex induces a large number of ASE in prostate cancer cells. To further confirm our observations, we conducted a thorough validation of predicted splice events using real-time PCR in three different prostate cancer cell lines, including androgen-sensitive LNCaP, castrate-resistant 22RV1, and metastatic PC3 cell lines. Together, our work provides strong evidence for a new role for AR-signaling in modulating ASE.

Interestingly, our work revealed that inhibiting AR-signaling using Casodex and enzalutamide significantly increases intron retention in comparison to DHT treatment. Intron retention is known to generate abnormal transcripts that are translated into immunogenic peptides, loaded on MHC-1, and presented to the immune system^50^. Therefore, patients with advanced-stage prostate cancer undergoing treatment with AR-inhibitors may have a higher neoepitope load and hence benefit from immune checkpoint inhibitors. Further studies will be necessary to predict and validate the immunogenicity of neoepitopes generated in response to AR-inhibitors, including identification of T cells infiltrating prostate tumors specific to predicted neoepitopes.

AR molecules in the cytoplasm dimerize and translocate to the nucleus in response to androgens and regulate the expression of target genes. Therefore, a majority of published reports have performed enrichment analysis with differentially expressed genes to understand the physiological significance of AR-signaling. In our work, we tested whether the AR-signaling regulated alternative splicing of pre-mRNA, has a different physiological role than that from differentially expressed genes. We found that while inhibiting AR-signaling led to expression changes of genes involved in EMT, it also led to splicing changes in pre-mRNA of genes regulating gene expression, including nucleic acid & protein transport, mRNA splicing, and proteasomal degradation. Because alternatively spliced genes were not differentially expressed in our analysis, we were able to discover a new physiological impact of modulating AR-signaling in prostate cancer cells. Besides, the role of AR-regulated splicing is further signified by our analysis showing that the majority of splicing occurred in characterized functional domains, including the transmembrane domain, coiled region, topo domain, metal binding, zinc finger binding, and activation sites.

The results in this work suggest that AR agonists and antagonists can dysregulate alternative splicing within the pre-mRNA of genes that regulate tumor biology. Because pharmacological modulators may have non-specific activity, it is possible that the observed ASE observed in response to treatment with AR-modulators is not mediated through AR. Employing RT-PCR, we demonstrate that inhibition of AR-signaling using pharmacological inhibitor or siRNA had a similar effect on the splicing of genes involved in cancer, including *AAK1, SYNE4*, and *MAN1A1*. Globally, we found a significant overlap as well as unique ASE events induced in different prostate cancer cells treated by pharmacological inhibitor or the genomic inhibition of AR. Because our global splicing analysis employed multiple cell lines to test this hypothesis, some of the overlap or unique ASE events identified could be because of cell line differences and not the effect of treatment. Therefore, the global comparison of ASE between the pharmacological inhibitor and genomic inhibition of AR signaling in prostate cancer cell lines needs further validation. Also, we cannot completely rule out the possibility of non-specific pathways being engaged by pharmacological inhibitors of AR to dysregulate splicing. However, we believe our analysis model and RT-PCR validation sufficient evidence to support our hypothesis that ASE events induced in response to treatment with AR inhibitor are in parts driven by direct modulation of AR expression.

The regulation of alternative splicing is primarily mediated by RBPs that interact with sequences flanking exon and introns as splicing enhancers or silencers, depending on the regulator and binding context. AR is a transcription factor; therefore, we argued that one mechanism through which it may regulate ASE is by modulating the expression of RBPs. We tested 112 well-characterized RBPs and found that only ESRP1/2 were transcriptionally regulated by pharmacological inhibitors of AR. Furthermore, consistent with the known mechanism for ESRPs driving splicing, we found that the binding sites for ESRPs were enriched or under-represented in the region within and flanking the AR-regulated cassette exon events. These data provide strong evidence for AR-ESRP axis driven cassette exons events in prostate cancer cells. We will need to perform ChIP-Seq to directly validate whether androgens or anti-androgens dysregulate ESRP binding around AR-driven cassette exons. Also, it would be interesting to study whether AR and ESRP bind within the same region and are part of a protein complex regulating splicing.

Because ESRP plays a critical role in EMT and tumor invasion in several tumor types^43^, we hypothesized that the AR-ESRP axis might be critical for promoting metastasis either by promoting EMT or promoting invasion. Moreover, although pharmacological inhibitors of AR-signaling suppress tumor growth, we and others have shown using *in vitro* and *in vivo* approaches that it also promotes invasion of tumor cells^44^. Hence, it is critical to identify new therapeutic targets that may alleviate accidental invasion promoting effects of inhibiting AR-signaling in advanced state prostate cancer. Our *in vitro* invasion assay found that silencing ESRP protein abrogates a Casodex mediated increase in the invasion of prostate cancer cell lines. Thus, a small molecule inhibitor for ESRP may serve as an attractive avenue to counter the invasion promoting phenotype of AR inhibitors.

Our work using prostate cancer cell lines showed that AR-signaling dysregulates the splicing of functionally relevant genes. Because AR-signaling is a hallmark of prostate cancer progression, we also investigated whether AR-regulated ASE is associated with the progression of disease. Our analysis showed that significant splicing events are occurring during different stages of prostate cancer progression. Contrary to an earlier study^45^, which had found that dysregulation of the expression of AR was associated with all stages of prostate cancer, we found an additional nine splicing events. Therefore, our work reveals a new potential area of inquiry into the underlying biology of prostate cancer initiation and progression to the castrate-resistant stage.

Furthermore, the transcriptomic analysis had only identified 15 genes that were differentially expressed between advPC and CRPC, thus making these two stages of cancer challenging to differentiate genetically. By focusing on alternative splicing, we have now identified an additional 13 genes that are differentially spliced between advPC and CRPC. Thus, a combined gene expression and splicing panel could potentially have a higher diagnostic value to discriminate patients in an advanced stage from the one with the castrate-resistant disease.

Lastly, we investigated whether the treatment of prostate cancer cells with pharmacological inhibitor enzalutamide may lead to an inadvertent switch in the splicing of a pre-mRNA and promote tumor progression. Our analysis found that dysregulation of AR-signaling in prostate cancer driven by enzalutamide treatment increases the inclusion of exon-2 of the *PLA2G2A* gene in prostate cancer cells. The translational significance of the inclusion of exon-2 was validated in the TCGA PRAD, where we revealed that the PLA2G2A isoform that includes exon-2 is a prognostic factor for outcomes (O.S), providing strong evidence for developing a therapeutic strategy that can mitigate the inadvertent pro-tumorigenic effects of inhibiting AR-signaling.

In conclusion, this study highlights the so-far undescribed role of AR in modulating gene expression via alternative splicing in prostate cancer. This discovery also opens a new therapeutic path and supports the rationale for using ESRP inhibitors in combination with AR-antagonists for the treatment of advanced-stage prostate cancer to counteract the AR-antagonist driven invasive phenotype.

## Materials & Methods

### Reagents

5α-Dihydrotestosterone (DHT), Casodex, and Enzalutamide were obtained from Sigma and were resuspended in DMSO (Sigma). Primers were designed manually and were purchased from Integrated DNA Technologies. Anti-ESRP1 (#21045-111-AP) & anti-ESRP2 (#23317-1-AP) were obtained from the Proteintech. E-Cadherin (4A2) (#14472) and Vimentin (D21He) (#5741) were obtained from the CellSignaling, while anti-aTubulin antibody (#A01410) was obtained from Genscript. All other reagents if not specified were obtained from Thermofisher Scientific.

### Cell Culture

The human cell lines LNCaP (androgen sensitive human prostate adenocarcinoma cells) and 22RV1 (human prostate carcinoma epithelial cell line derived from a xenograft that was serially propagated in mice after castration-induced regression and relapse of the parental, androgen-dependent CWR22 xenograft) were obtained from ATCC and were cultured in RPMI 1640 medium. PC3 (metastatic prostate cancer cells isolated from bones, ATCC) were maintained in F-12K medium. All culture medium was supplemented with 10% HyClone Defined Fetal Bovine Serum (GE Healthcare) and 1% Pen/Strep (Invitrogen) unless specified. Cell cultures were tested every 6 months for cross-contamination using human STR profiling cell authentication service provided by ATCC. Mycoplasma contamination was tested using MycoAlert Mycoplasma Detection Kit (Lonza).

### Western Blotting

Cells were dissolved in RIPA buffer (sigma). Protein concentration was measured by BCA protein assay reagent kit (Pierce, Rockford, IL, USA), as described previously. Proteins were fractionated on 10% SDS-PAGE, and transferred by electrophoresis to nitrocellulose transfer membrane (GE). Membranes were incubated with primary antibodies for overnight. Horseradish peroxidase-conjugated antibody anti-mouse IgG and anti-rabbit IgG (Dako) were used to detect immunoreactive bands and binding was revealed using enhanced chemiluminescence (Pierce). The blots were then stripped and used for further blotting for control antibody. Unless otherwise specified, displayed western blots are representative images of at least 2 independent experiments.

### siRNA Transfection

PC3, LNCaP, and 22RV1 cells were transfected with siRNAs targeting ESRP1/ESRP2 kinases and negative control (SiGenome ESPR1 #D-020672-01, On-Target ESPR2 #J-014523-05, Darmacon, and Silencer Negative control, #4390843 Thermofisher). Cancer cells were seeded in a 6-well plate. 200nM siRNA/well were used for transfection using 5µl/well of Hi-Perfect (Qiagen) following manufacturer’s recommendation.

### 2D Invasion Assay

Reduced growth factors Matrigel (BD Biosciences) was polymerized in 8 mm pore cell inserts (Sarstedt) prior to the addition of cells. LNCaP and 22RV1 (5 × 106 cells) were seeded into the insert containing Matrigel in serum free media. 20% FBS medium were used as an attractant in the bottom chamber, and cells were allowed to migrate for additional 24hr. The inserts were removed, and migrated cells were fixed with 4% paraformaldehyde and stained with crystal violet. The inserts were then imaged, and migrated cells were counted, hence, providing a quantitative value for migrated cells across the membrane.

### Human Transcriptomic Array

LNCaP cells were cultured for 3 days in the RPMI-1640 with 10% Charcoal-Stripped Fetal Bovine Serum and treated with either 10nM DHT, 10mM Casodex, or DMSO for 24 hours. Total RNA was extracted using the RNeasy mini kit (Qiagen) and quantified using Nanodrop ND-100 Spectrophotometer (Thermo Fisher Scientific). The quality of RNA was assessed using Agilent 2100 Bioanalyzer (Agilent). Biotinylated cDNAs were prepared from a minimal 100ng of total RNA using Life Technologies WT-plus RNA Amplification system (Ambion). Following the amplification, cDNA was fragmented, hybridized on Affymetrix GeneChip Human Transcriptome Array 2.0 (HTA2.0) chips and non-specific bindings was washed as per manufacturer’s recommendations. The fluorescence intensity of the arrays was scanned using the Affymetrix Scanner and the raw data. The raw data were analyzed using the Transcriptome Console Software (TAC 2.0), which allows for the identification of differentially expressed genes and leverages information from the junction and exons probes to detect alternative splicing events and possible transcript isoforms that may exist in samples.

For microarray data analysis, two parallel analysis (gene-level and alternative-splicing level) were performed using HTA 2.0. Data were normalized using quantile normalization and the background was detected using the built-in detection above background algorithm (DABG). Only the probesets that were characterized by a DABG *p-value* < 0.05 in at least 50% of the samples were considered as statistically significant. Genes were considered to be differentially expressed when Fold Change (FC), log <= −2.0 or >= +2.0 and FDR corrected *p*-value <= 0.05. The splicing level analysis as also carried out using TAC 2.0 software, which determines the Splicing Index (SI) of a gene. The SI corresponds to a comparison of gene-normalized exon-intensity values between the two analyzed experimental conditions (*). Additional criteria used besides SI: FDR corrected *p*-value £ 0.05, a gene is expressed in both conditions tested, a Probeset Ration (PSR)/Junction must be expressed in at least one condition, a gene must contain at least one PSR value, and a gene cannot be differentially expressed.

### Reverse Transcription PCR Validation for Splicing Events

A total validation of 15 splicing events was performed on three prostate cancer cell lines including LNCaP, 22RV1, and PC3 at various disease state (primary prostate cancer, castrate-resistant, and metastatic prostate cancer). Briefly, the prostate cancer cells were cultured for 3 days in 10% CSFBS and were treated with either DMSO, 10nM DHT, or 10mM Casodex for 24 hours. Total RNA was extracted, and cDNA was prepared using SuperScript IV First-Strand Synthesis System (Thermo Fisher Scientific). cDNA samples were amplified using SYBR® Green PCR Master Mix on the Applied Biosystems 7500 Detection System. Spicing-specific primers included two primer pairs, one for monitoring the expression of constitutive exons within all the isoforms of a gene and another for measuring changes in the alternatively spliced region. Furthermore, specificity and efficiency for primers were analyzed by running RT-PCR with series of cDNA dilutions and specific amplification for every assay were confirmed by melt curve analysis. The amplified transcripts were quantified using the comparative ΔΔCt method. GAPDH and HPRT were used as the internal control. Splice Index (SI) was calculated for (A) by normalizing fold change (FC) to the average FC of (C) for each splicing event. All assays were run in triplicates and were repeated 3 times. A raw Ct of 35 is used as the limit of detection: Ct values are set at 35 for any replicates with Ct values not determined or >35.

### Raw data processing, alignment analysis, and identification of differentially expressed genes and alternative splicing events

High-quality RNA samples were extracted and illumine library were constructed as described earlier. Libraries were pooled and diluted for sequencing with a 1% PhiX spike-in according to Illumina protocol. The pool of library was loaded onto the HiSeq was performed using a 300-cycle high output v2 kit. Reads were obtained from sequencer or were downloaded from GEO. Adapter sequences and invalid reads containing poly-N and low-quality were removed using the FastX tool kit (v 0.0.14). The quality of reads was then confirmed using fastqc tool kit (v 0.11.5). All downstream analysis used the cleaned reads. The clean reads were mapped to the ENSEMBL built GRC38 using the STAR aligner (v2.5.3a) using ENCODE option as described in the STAR manual. Differentially expression of genes was obtained using the DESeq2 method as described earlier. Subsequently, we used rMATS (version3.0.8) to identify differentially ASE between the two sample groups. Briefly, rMATS uses a modified version of the generalized linear mixed model to detect differentially ASE from RNA-Seq data with replicates, while controlling for changes in expression at gene-level. In addition, it also accounts for exon-specific sequencing coverage in individual samples as well as variation in exon splicing levels among replicates. rMATS was run using the default parameters and then significant splicing events were filtered using a stringent cut-off of FDR ≤ 0.05and deltaPSI ≥ 10%. The R package, Maser, was used for extracting and visualizing splice events. and then significant events were extracted and further analyzed using the R package.

### Motif Enrichment Analysis

We employed rMAPS2 analysis to determine the binding patterns of splicing factors and RNA binding proteins within significantly detected exon skipping ASE between two treatment groups. We collected well-characterized 115 known binding sites of RNA binding proteins. For each motif, the analysis scanned for motif occurrences separately in exons or their 250bp upstream or downstream intronic sequences. Furthermore, for the intronic sequences our analysis excluded the 20bp sequences within the 3’ splice site and the 6bp sequences within the 5’ splice site. In addition, by default the alternative exons without splicing changes as defined by rMATS FDR > 50%, maxPSI > 15%, and minPSI < 85% were treated as control exons. For each motif tested, the analysis counted the number of occurrences and motif score in the upstream exon, upstream flanking intron, target exon, downstream flanking intron and downstream exon separately. The p-value for motif enrichment was calculated using the Wilcoxon’s rank sum test for each sliding window between upregulated versus control or downregulated versus control exons.

### Functional Annotation of DEGs and ASE

The ReactomePA & the clusterProfiler were used to generate lists of the Gene Ontology (GO) terms enriched in the DEGs and ASE. The integration of protein features to splicing events was carried out using maser package. Briefly, maser enables systematic mapping of protein annotation from UniprotKB to splicing events and determine whether the splicing is affection regions of interest containing known domains or motifs, mutations, post-translational modification and other described protein structural features.

## FIGURE LEGEND

**Supplementary Figure 1–3:**
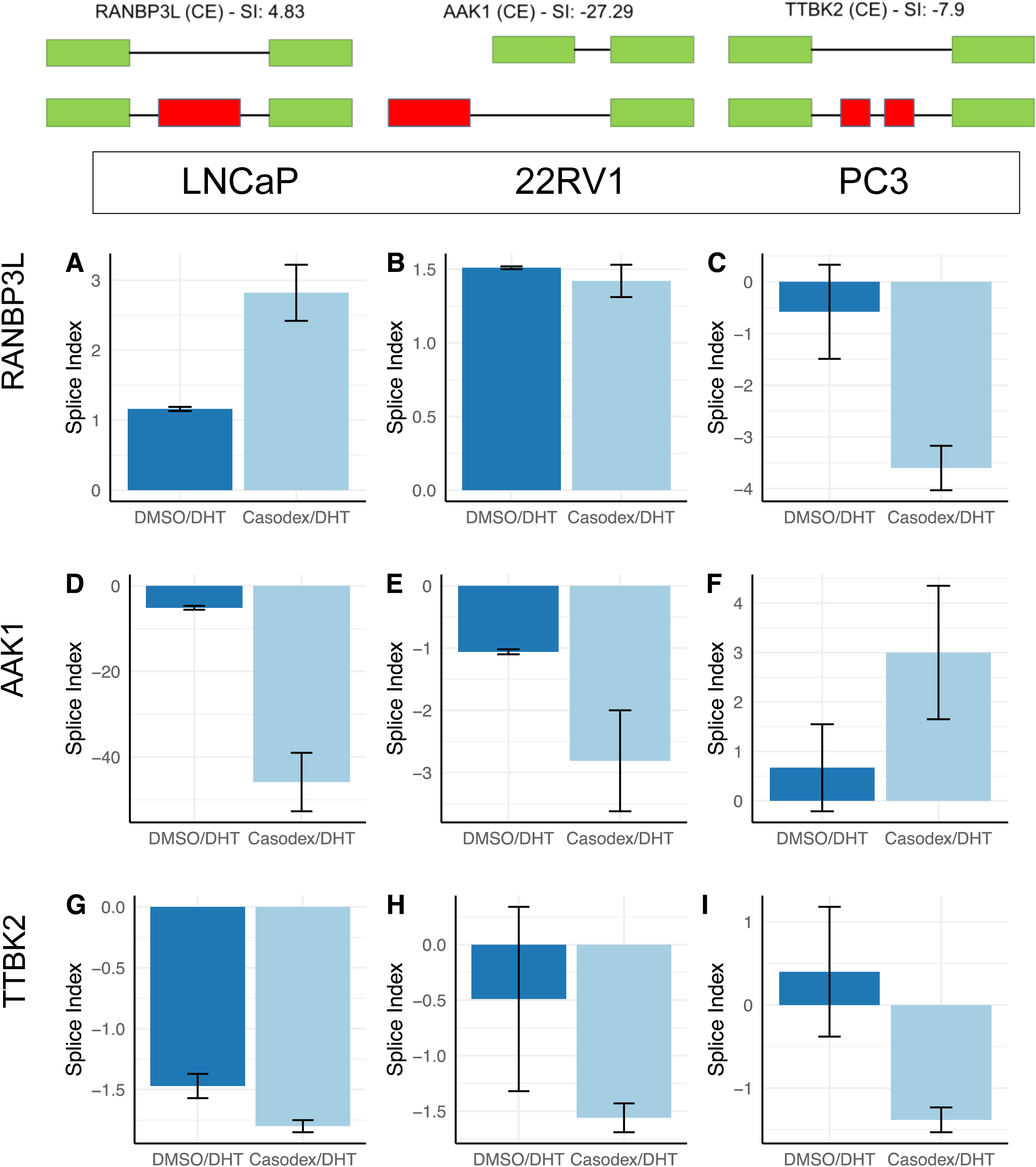
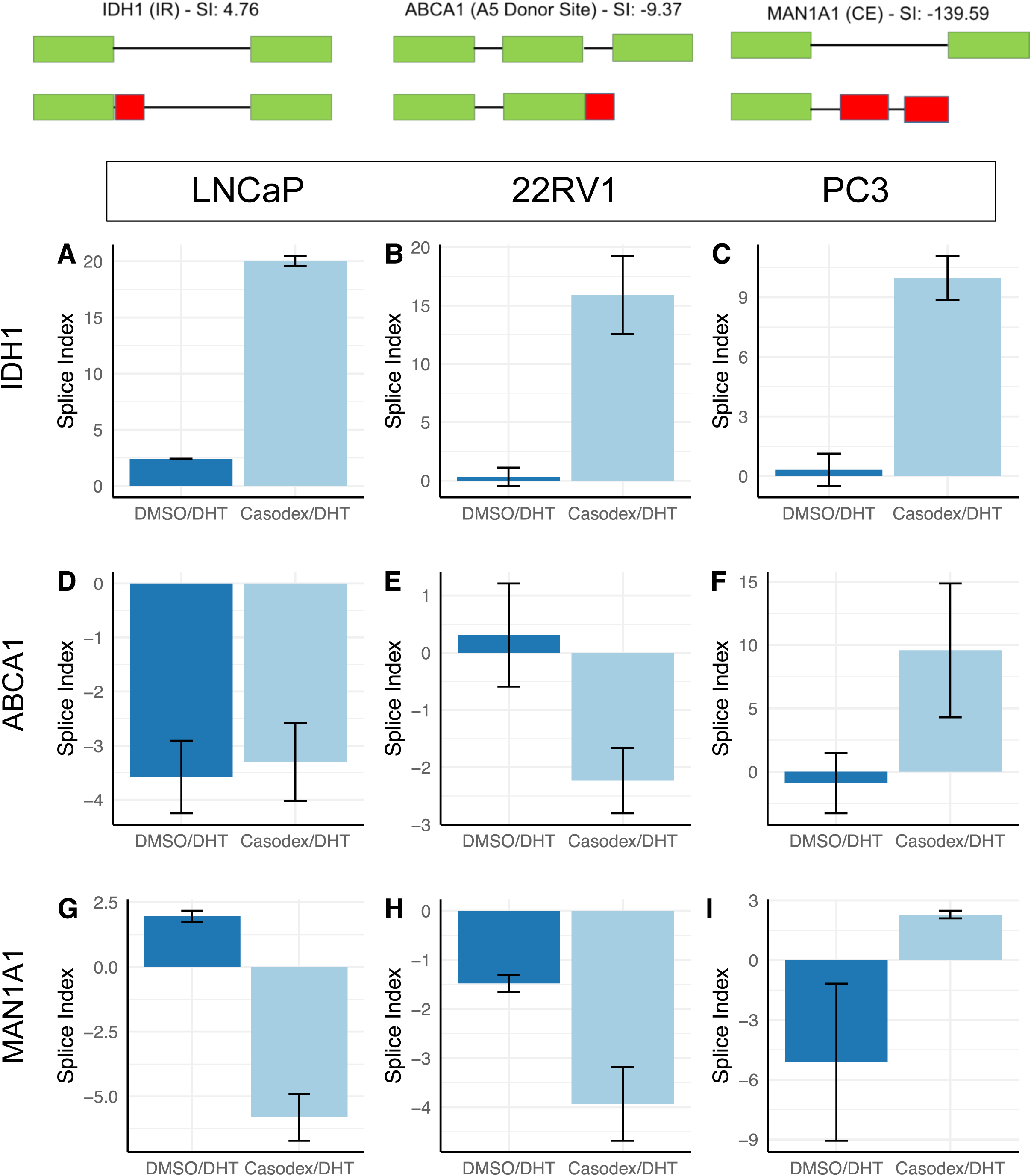
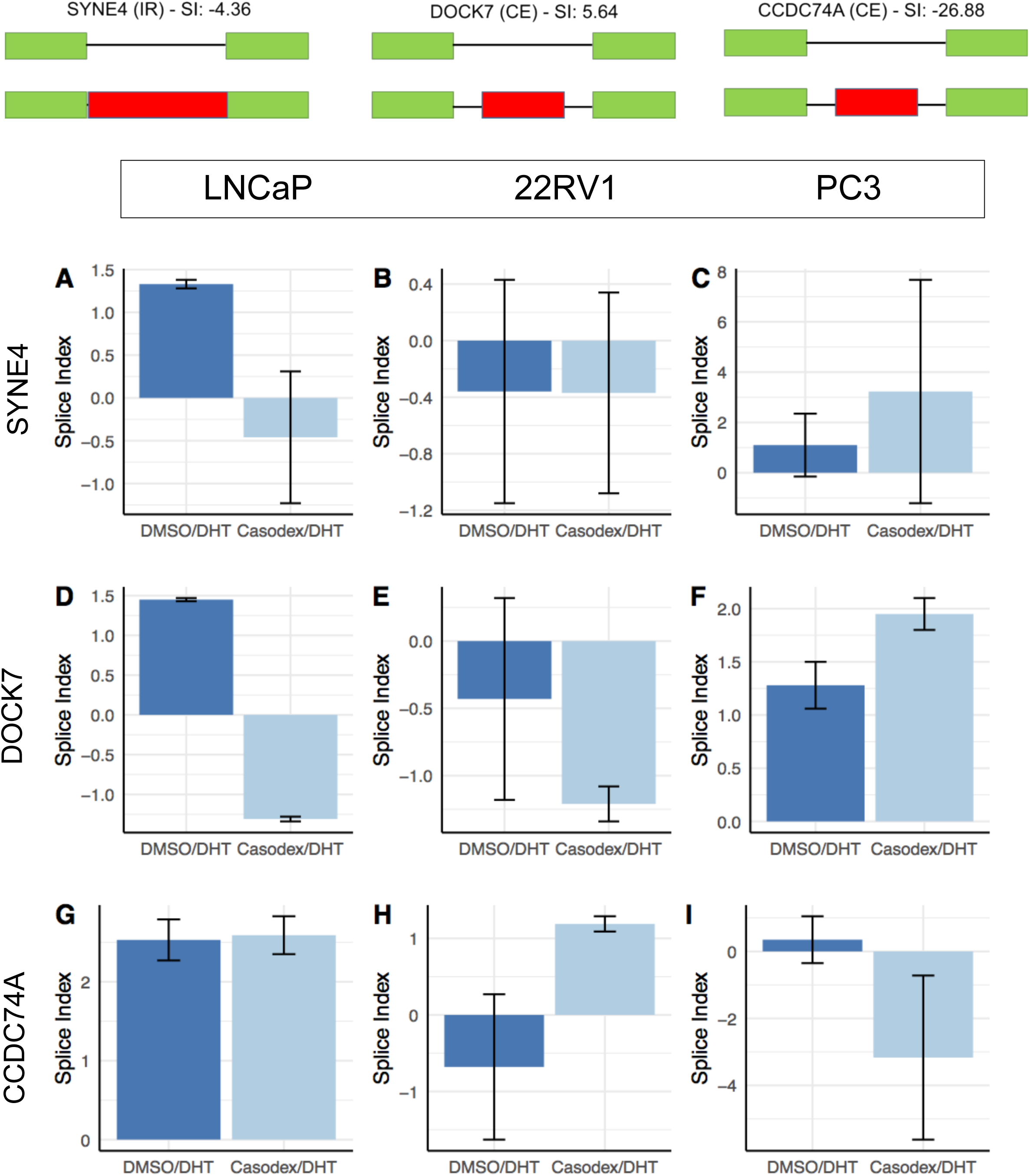
RT-PCR validation of alternative splicing events predicted using splicing array. Bar-graph showing the average splice index of genes including RANBP3L, AAK1, TTBK2, IDH1, ABCA1, MAN1A1, SYNE4, DOCK7, and CCDC74A across LNCaP, 22RV1, and PC3 cells from RT-PCR data calculated using the ΔΔCt method relative to an endogenous reference gene (HPRT or GAPDH).

**Supplementary Figure 4:**
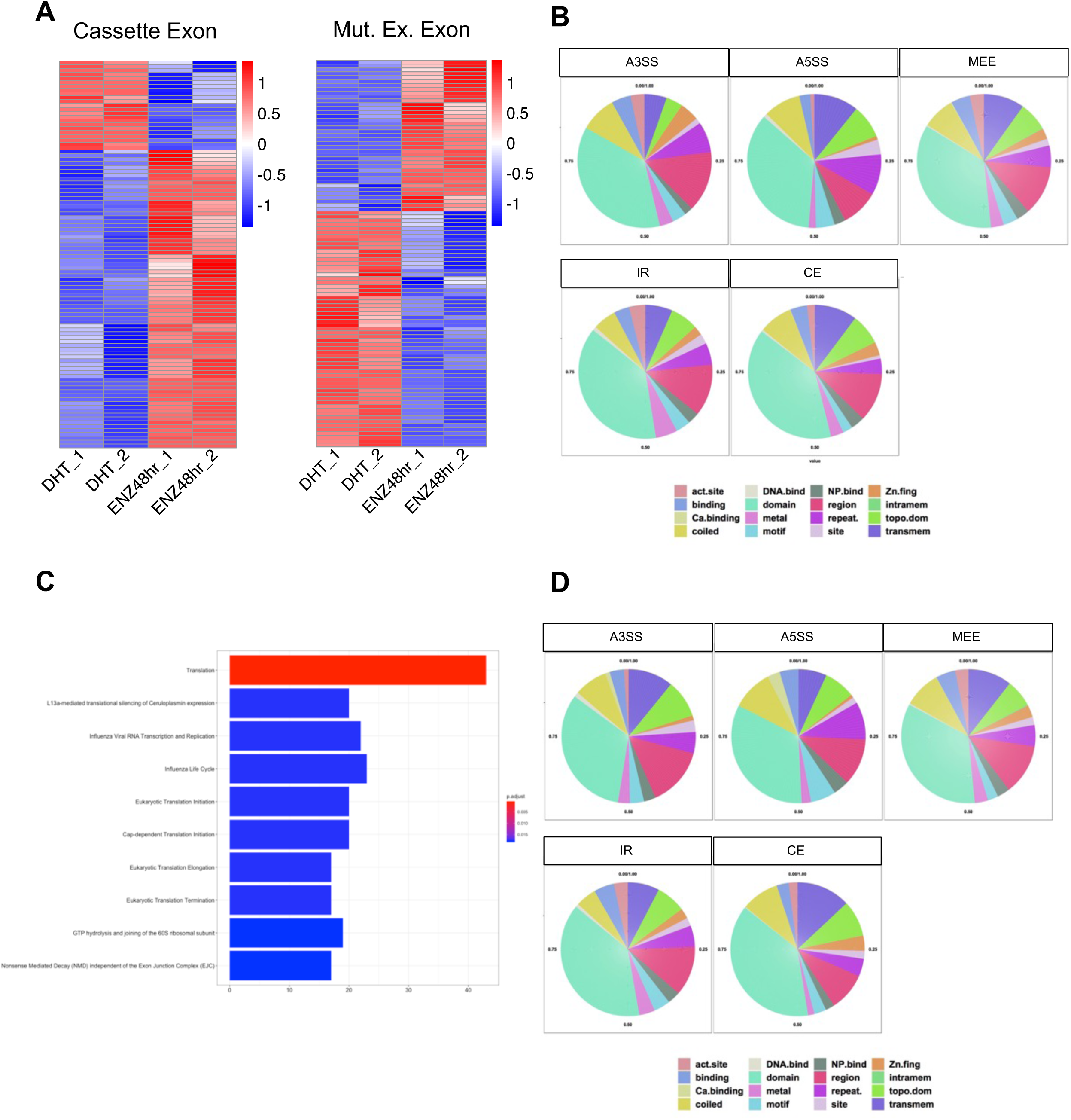
(A) The heatmap comparing showing top–100 significantly different PSI for MEE and CE across different samples including LNCaP cells treated with DHT or enzalutamide. (B) Pie-charts show the frequency of splicing events induced by enzalutamide in comparison to DHT treatment of LNCaP cells that are mapping to the protein functional domains including transmembrane domain, coiled region, topo domain, metal binding, zinc finger binding, activation site for protein, and others. (C) Bar-graph showing the GO pathways enriched for a common set of genes undergoing alternatively splicing in response to treatment with DHT, enzalutamide or siRNA against AR. (D) Pie-charts show the frequency of splicing events induced by the genomic inhibition of AR in MDA-PCa-2B cells that are mapping to the functional protein domain.

**Supplementary Figure 5:**
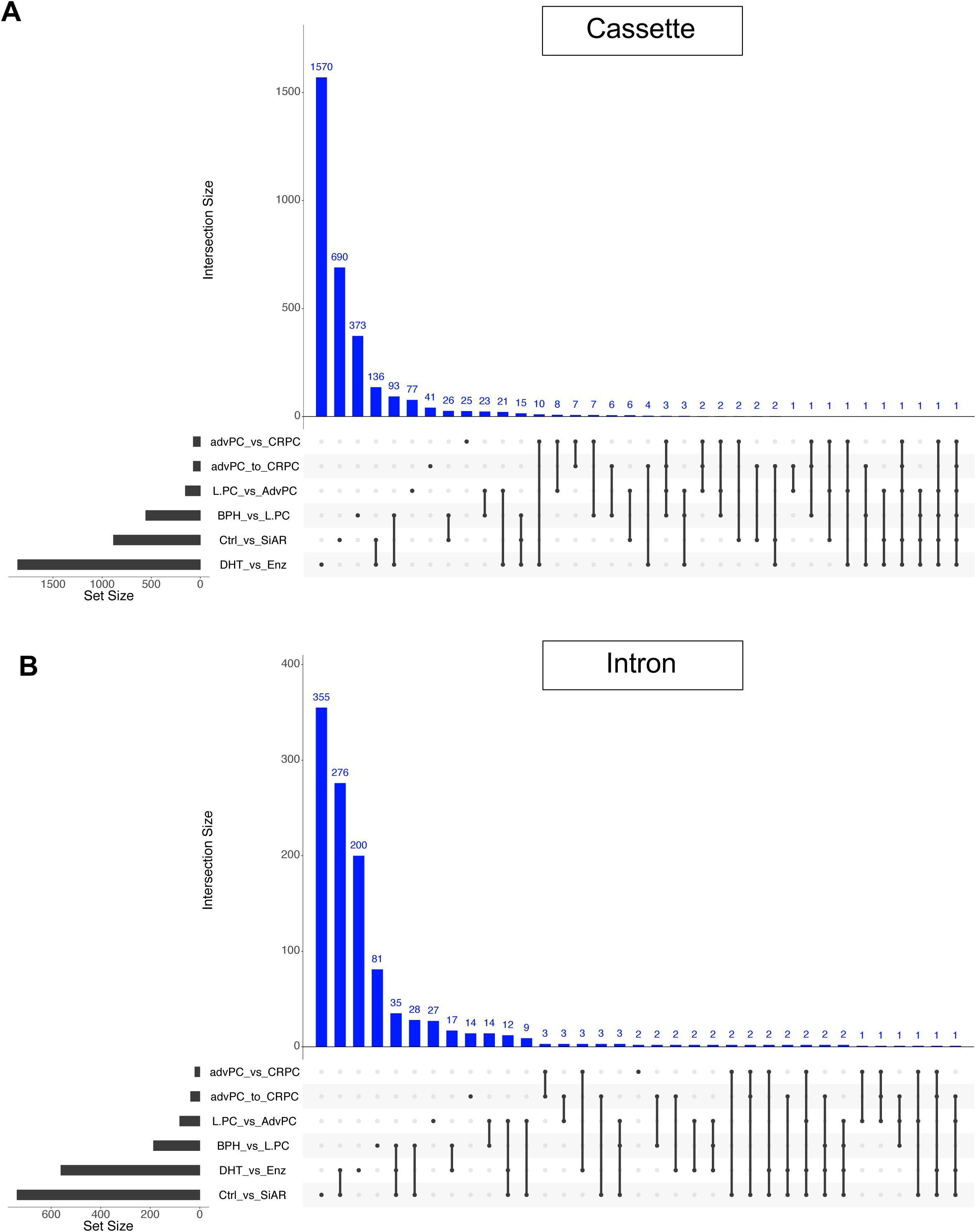
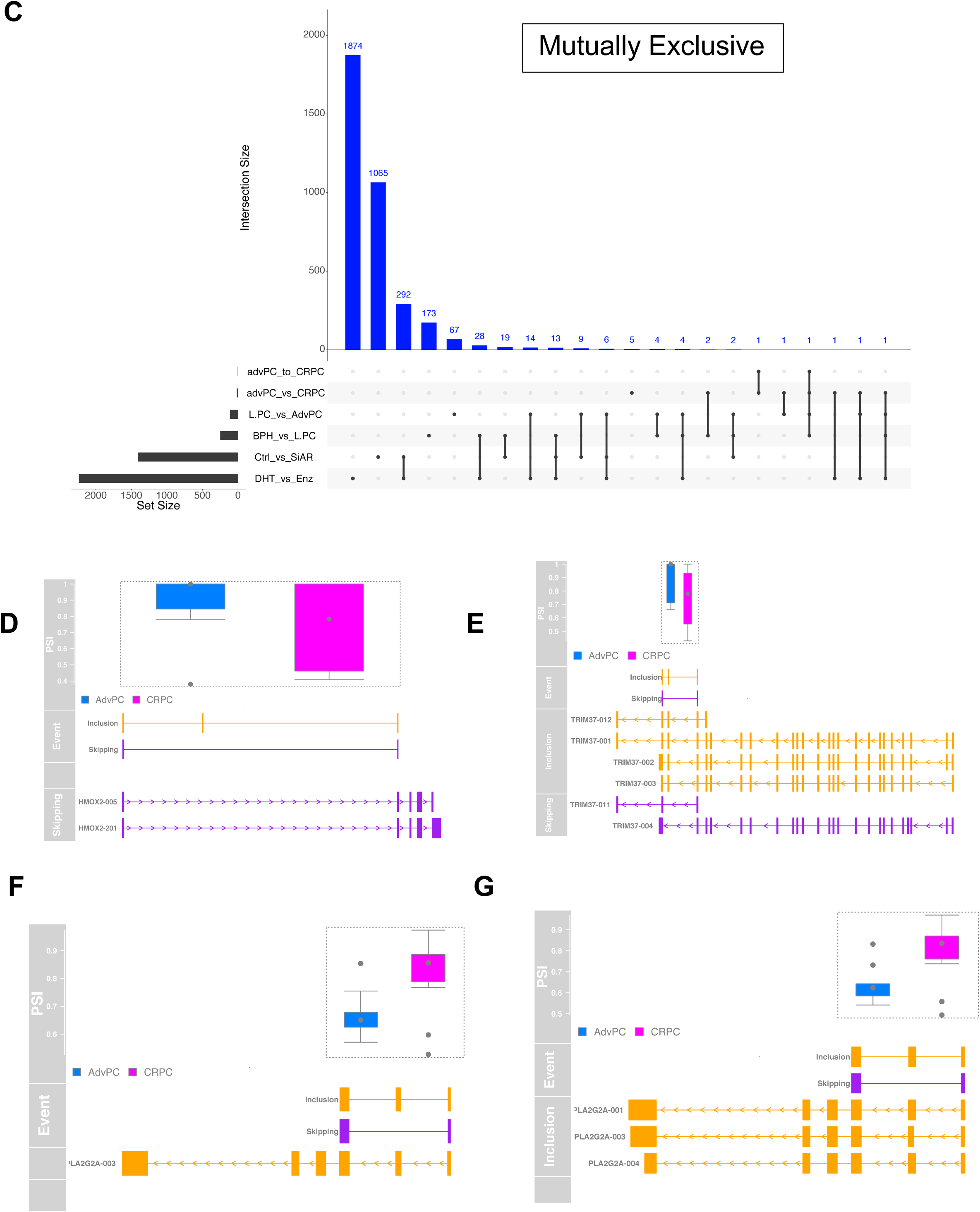
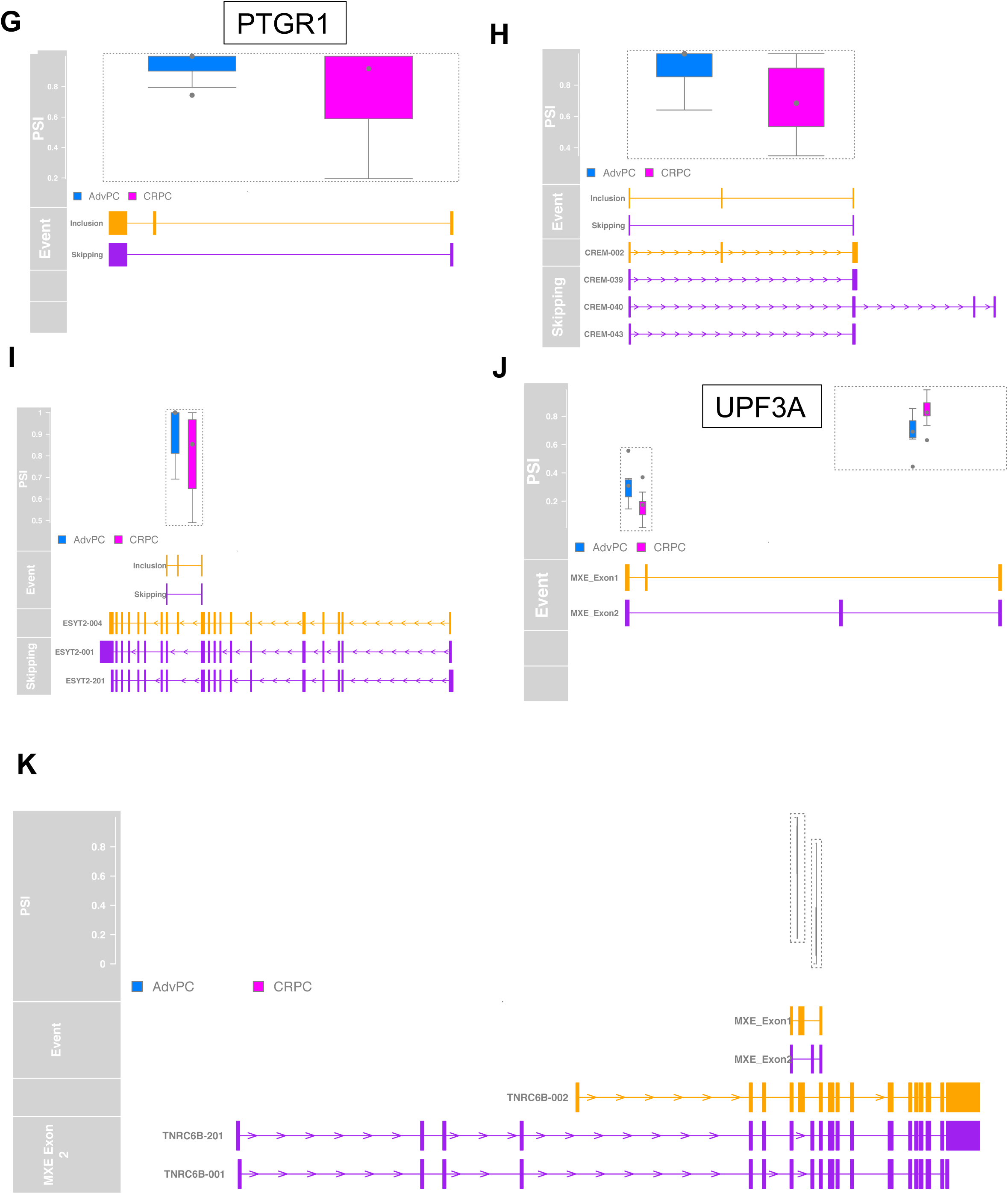
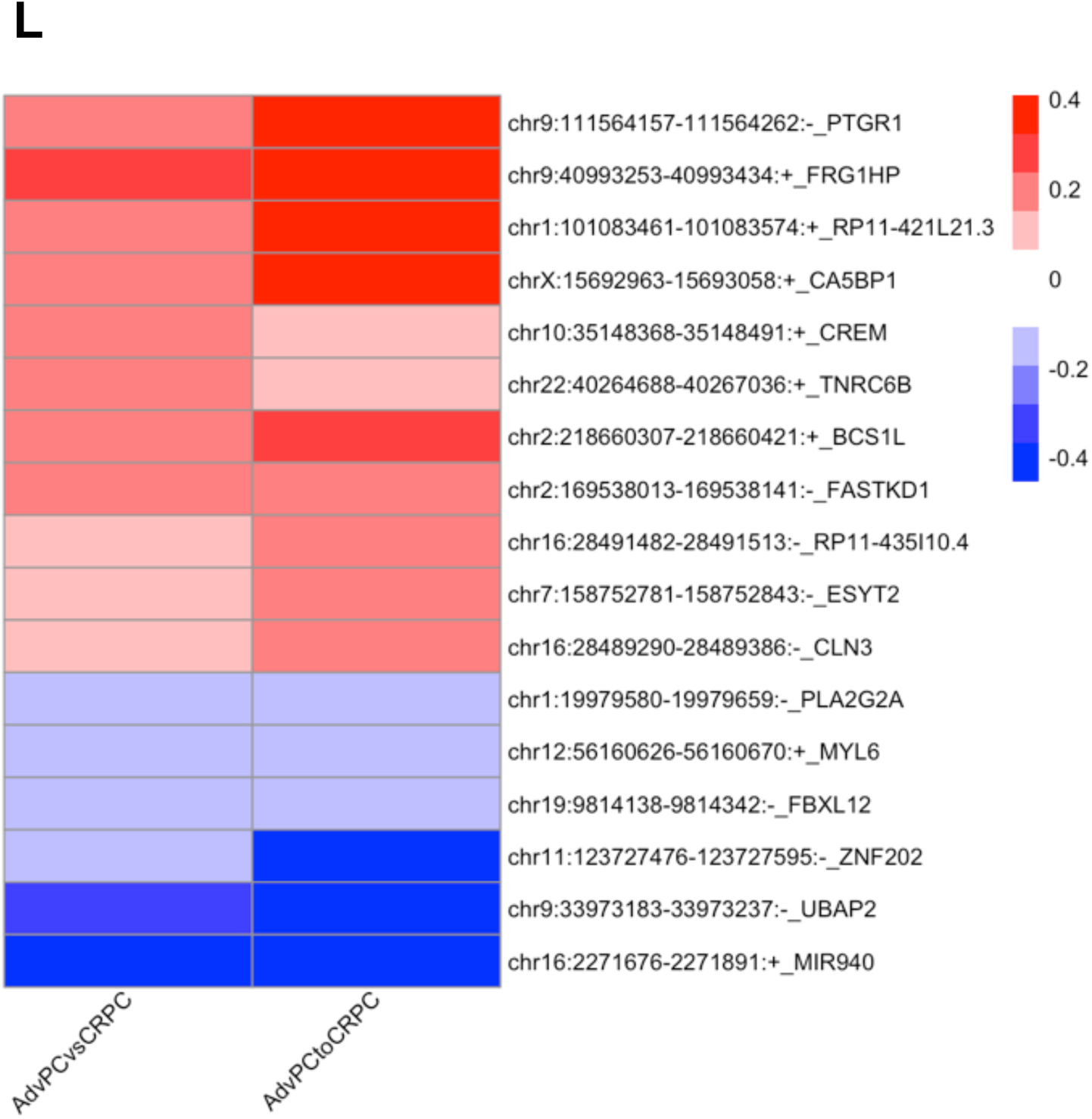
(A-C) Upset plot showing an overlap and unique cassette exon, Intron retention, and mutually exclusive events across comparison groups including patients with BPH, L.PC, AdvPC, CRPC, LNCaP cells treated including DHT or enzalutamide, MDA-PCa-2b cells treated with scrambled siRNA or siRNA. (D-K) A combined transcript plot including a box plot comparing the PSI between patients with AdvPC and CRPC, the schematic of the splicing event, and predicted ensembl transcript plot for cancer-relevant genes. (L) Heatmap comparing the PSI for the ASE predicted to be common across AdvPC vs. CRPC and a longitudinal comparison within patients who progressed from AdvPC to CRPC.

**Supplementary Figure 6:**
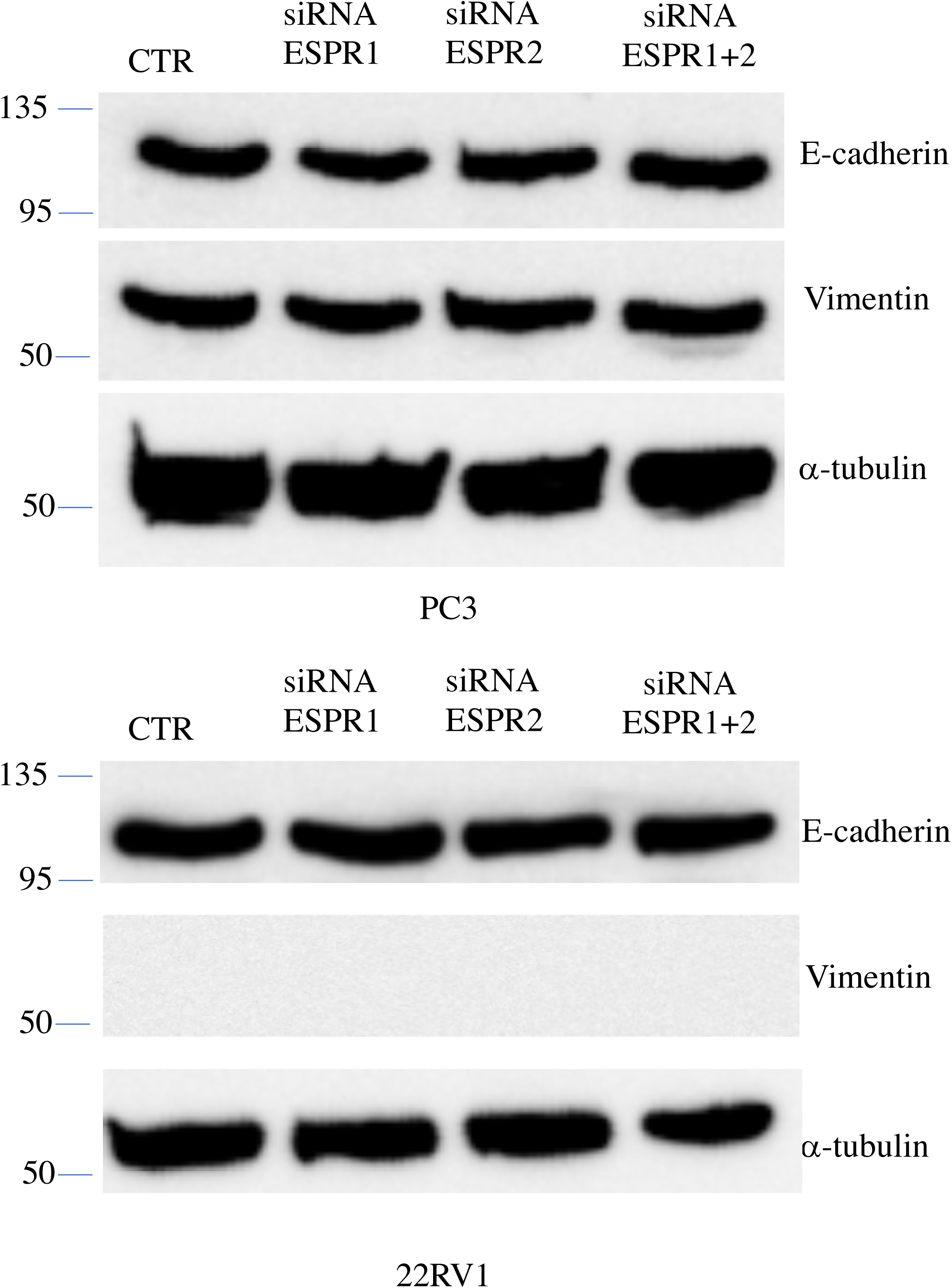
PC3 and 22 RVI cells were transfected with indicated siRNA, after 24 h, western blotting of EMT markers (E-cadherin, and Vimentin) was performed using Tubulin as loading control.

## Acknowledgements

This work was supported by grants from the intramural research program of the Division of Cancer Epidemiology and Genetics, National Cancer Institute, National Institutes of Health. This work utilized the computational resources of the NIH HPC Biowulf cluster and DCEG CCAD cluster. (http://hpc.nih.gov)

## Competing interest

The authors declare no competing financial interests.

